# Dengue Virus Alters Sialic Acid Residues Configuration in Macrophages

**DOI:** 10.1101/2021.02.26.433067

**Authors:** Javier Serrato-Salas, Isabel Cruz-Zazueta, José Luis Montiel-Hernández, Judith González-Christen

## Abstract

The activation of the innate immune response requires sialic acid residues removal. Nevertheless, it is unknown the role for these changes during the Dengue virus infection. We determine if during Dengue virus infection, the sialic acid residues alter on the macrophages. The human monocytic cell line THP-1 was differentiated into macrophages and were infected with Dengue virus. The changes in sialic acid were evaluated by lectin blot in the cellular lysate. The activity of neuraminidase was defined by RT-PCR and fluorescence assays. Macrophages infection with DENV-2 reduces α-2,6 sialic acid residues at 24 h, and α-2,3 sialic acid residues lower at 48 h in some proteins. Transcriptional profile and enzymatic activities of Neu-1 showed a narrow decrease. Sialic acid residues oscillation in varied conformations and times suggest it a role of a selective mechanism to remove these residues. The lesser participation of Neu-1 in this process could be concomitant to other similar enzymes such as sialyl-transferases, or the phenomenon requires minimal activity to have a relevant biological function.

## Introduction

Dengue virus (DENV) is the etiological agent of the disease-bearing a wide spectrum of clinical outcomes, from asymptomatic until severe complications that may be life-threatening. (Bhatt et al. 2013; Leta et al. 2018). A new specific antiviral treatment acknowledges getting a deeper understanding of the earlier stages in cell infection.

### The role for sialic acid in innate immune response activation

The viral replication cycle begins with the mature virion envelope and capsid binding to receptors outside the cell, and further entering the cell. Then, viral molecular patterns are recognized by host receptors of the innate immune system to mount a biological response to contain potential damage. (De La Cruz Hernández et al. 2016; Chaichana et al. 2014; De La Cruz Hernández et al. 2014).

As general, viral genomic templates, as single-strand RNA uncoiled poly (A), double-strand RNA in the cytoplasm, as well as unmethylated DNA, are censored by TLR receptors −3, −7 and, −9, respectively, which dimerize and activate a subsequent signaling cascade. (Rothman 2011, 2009; Srikiatkhachorn et al. 2017; Rodenhuis-Zybert et al. 2010; Georgel et al. 2009).

Moreover TLR activation, sialic acid takes part in other processes as adhesion and leucocyte leakage, cell to cell communication, among others. (Almaraz et al. 2012; G.-Y. Chen et al. 2014; Glanz et al. 2019; Abdulkhalek et al. 2011).

LPS binding to TLR-4 rises neuraminidase activity of Neu-1 and Neu-3 in dendritic cells and monocytes. In human macrophages, LPS binding to TLR-2, −3, and −4 induces neuraminidase activity of Neu-1, which hydrolyzes α-2,3 residues in the β-galactoses present in glycoproteins, this later induces the receptor dimerization and downstream signaling activity. (Bonardi et al. 2014; Amith et al. 2010).

Neu-1 removes α-2,3 residues in galactose from receptors TLR-4 and −2, allowing the dimerization and activation alongside the metalloprotease activation MM-9. The neuraminidase activity in dendritic cells-derived monocytes increments the secretion of IL-6, IL-12p40, and TNF induced by LPS, neuraminidase activity inhibition using oseltamivir phosphate, declines cytokine production, as a response to LPS stimuli. The neuraminidase activity in LPS-stimulated cells enhances phosphorylation of transcriptional factor NF-κBp65, which favors the transcription of proinflammatory cytokines. This phenomenon has been detailed before for TLR-7 and −9. (Abdulkhalek et al. 2013, 2011).

It has been proven Neu-1 translocation in vesicles towards cell surface joined to MHC-II molecules. (Liang et al. 2006). Phagocytosis could be regulated similarly for Neu-1 activity. (Seyrantepe et al. 2010).

### The role for sialic acid in Dengue virus infection

In mosquitos, it has been reported DENV recognizes α-2,6-sialic acid sites over the cell surface, which facilitates virion entry. (Cime-Castillo et al. 2015; Conway et al. 2014; Thaisomboonsuk et al. 2005).

In human, lack of glycosylated residues in domain III of viral envelope protein does not allow membranes fusion and further viral propagation. (Cruz-Oliveira et al. 2015; De La Guardia et al. 2014).

NS1 acts as a PAMP and activates murine macrophages and mononuclear cells derived from human peripheric blood through TLR-4 receptors leading to an inflammatory response and cytokine and chemokine releases. NS1 recombinant treatment disturbs membrane integrity in a monolayer of endothelial cells, this phenomenon was inhibited with a TLR-4 antagonist. (Beatty et al. 2015; Glasner et al. 2017).

NS1 induces TLR-2 and −6 activation, TRL-6 prolonged activation resembles to be an important component in the immunopathogenesis. (J. Chen et al. 2015).

In addition to, NS1 increases cathepsin L activity, at the same time heightens and activates heparinase in endothelial cells degrading heparan sulfate, which may contribute to plasma leakage. (Puerta-Guardo et al. 2016).

In this work, it was evaluated the sialic acid changes on macrophages infected with Dengue virus serotype 2 and reveal if neuraminidase is the enzyme related to removing sialic acid residues.

## Methods

The project was realized from January 2016 to October 2017 in Innate Immunity Laboratory of the Pharmacy Faculty facilities in University Autonomous of Morelos State (FF-UAEM). THP-1 human monocyte cell line (ATCC TIB 202, USA) was used for *in vitro* assays, which has been demonstrated as being a robust model of monocytes/macrophages. Cells were grown in Advance RPMI 1640 media (GIBCO, Thermo-Fisher, USA) supplied with 3% fetal bovine serum (GIBCO, Thermo-Fisher, USA) in a CO_2_ incubator at 37 °C. To differentiate monocytes to macrophages it was added PMA 10nM as a final concentration (Sigma-Aldrich, USA) over 72 h. Dengue virus serotype 2 New Guinea C strain was propagated in C6/36 cells in the laboratory of Immunity and Infection of Center of Research for Infectious Diseases of the National Institute of Public Health (CISEI-INSP), according to available protocols described elsewhere. For infection assays, cells were washed twice with PBS, virus aliquot was diluted in no-serum media for a MOI of 1, then incubated at 37 °C for 2 h, after this supernatant was discarded and replaced with fresh media for all experimental groups.

### Macrophage infection test

An immunocytochemistry assay was performed using antibodies against envelop protein (Genetex GTX43295, USA) diluted 1:45 in PBS at 25 °C for 1 h. Secondary antibody anti-IgG mouse coupled with phycoerythrin (Biolegend, USA) was used 1:100 in PBS at 25 °C for 45 min. Pictures were taken with an epifluorescence microscopy Nikon (Japan). The images were processed using ND2 software (Nikon, Japan).

RT-PCR was realized as additional proof to detect viral genome presence. Total RNA was extracted from cells using TRIreagent (Thermo-Fisher, USA) following the manufacturer’s instructions. cDNA synthesis was performed with RevertAid First Strand cDNA synthesis (Thermo-Fisher, USA) using random hexamer primers. PCR was made using PCR Master Mix (Thermo-Fisher, USA) according to standard conditions, 35 cycles (94 °C for 20 s, 60 °C for 30 s, 72 °C for 1 min) and final extension 72°C for 5 min. Primers sequences to detect Dengue virus presence were, DENV ALL F 5’-CAATATGCTGAAACGAGAGAGAA -3’, DENV ALL RV 5’ - CCCCATCTATTCAGAATCCCTGC -3’. The amplicon from the first PCR was used for a nested PCR, diluting 1:200 and substituting reverse primer for DENV-2 RV 5’-TGCTGTTGGTGGGATTGTTA -3’ using the same PCR running conditions. PCR products were visualized in agarose gel using a DNA fluorescent intercalator Safe-Green (ABM, USA) through electrophoresis.

#### Sialic acid changes determination in infected macrophages

Macrophages were infected with DENV, as positive control cells were stimulated with LPS 10 ng/mL, and incubated during 24 and 48 h. The supernatant was discarded, cells were washed with PBS. For lectin blot assay, cells were differentiated in 25 cm^2^ culture flasks with a concentration of 1×10^6^ cells/mL. After treatment cells were lysed using RIPA buffer with proteases inhibitors. Protein separation was realized through SDS-PAGE. Proteins were transferred to a nitrocellulose membrane. Lectins used for the assays, SNA for α-2,6 residues (diluted 1:3 500), MAA for α-2,3 residues (diluted 1:1 000) and PNA for β-1,3-galactosamine residues (dilution 1:1 000). Lectins were incubated overnight at 4 °C. Revealing was carried out using streptavidin-peroxidase (Sigma-Aldrich, E886, USA) over 1 h, with Western Lightning Plus ECL (Perkin Elmer, USA). Images were recorded with ChemiDoc XRS (Bio-Rad, USA).

#### Neuraminidase activity in infected macrophages

Using the equal experimental conditions for differentiation and infection, neuraminidase-specific inhibitor DANA was added to a final concentration of 100 μM (N-Acetyl-2,3-dehydro-2-deoxyneuraminic acid, Merck, USA). It was used for 48 h prior stimulus.

Neuraminidase transcriptional activity was evaluated using inhibitor oseltamivir phosphate (Tamiflu, Roche, Switzerland) was added to a final concentration of 200 μM. RT-PCR transcript levels were determined in the laboratory of Infection and Immunity of CISEI-INSP. Primers sequences were Neu-1 Fwd 5’ - ACCTTGGGGCAGTAGTGAG - 3’ Neu-1 Rev 5’-TCCCGCTGTTTCTGAATACCA-3’ and β-actin Fwd 5’-GCTCCGGCATGTGCAA – 3’ y β-actin Rev 5’-AGGATCTTCATGAGGTAGT – 3’ as control.

Neuraminidase enzymatic activity was established at 24 h post-infection, cells have fasted for 3 h previous to the stimulus, enzymatic activity was revealed using 4-MUNANA substrate (Sigma-Aldrich, USA), which is hydrolyzed for neuraminidases and produces the free soluble fluorescent molecule 4-methyllumbelliferyl, with a range for fluorescence excitation/emission in 365nm/450nm wavelength. Images were captured 1 to 5 minutes after addition using an epifluorescence microscopy Zeiss (Germany) with the software Axio Imager, Zeiss (Germany).

## Results

### Sialic acid changes determination in infected macrophages

Macrophages were infected with MOI 1 according to laboratory protocol settled. Images showed the presence of the virus in cytoplasm and the presence of the viral genome for RT-PCR (Cross-ref).

After infection, cell lysate was obtained and changes in sialic acid were determined through SDS-PAGE with the lectin-blot described. The average result of the assays (Cross-ref).

Semiquantitative analysis was set up of the blot by pixel density. For SNA (α-2,6) in infected cells proteins weights between 100-250 KDa diminish at 24 h; proteins between 55-70 KDa decrease at 48 h. For MAA (α-2,3) in infected cells, sialic acid content diminishes at 24 h, and there were no significant changes at 48 h. For PNA (β-1,3-gal) in infected cells showed a reduction at 24 h, but none at 48 h (Cross-ref).

Changes for each lectin are highlighted in (Table 1 cross-ref).

**Table 1.** Resume the changes in sialic acid content for the three lectins used at 24 and 48 h. ↑ protein with an increase. ↓ protein with a decrease. S/C protein with no changes compared to control.

|  | SNA |  |  | MAA |  |  | PNA |  |
| --- | --- | --- | --- | --- | --- | --- | --- | --- |
|  |  |  |  | 24 h |  |  |  |  |
| Protein | DENV | LPS | Protein | DENV | LPS | Protein | DENV | LPS |
| 67 | ↓ | S/C | 116 | ↓ | S/C | 123 | ↓ | S/C |
| 59 | ↓ | S/C | 90 | ↓ | S/C | 91 | ↓ | S/C |
| 55 | ↓ | S/C | 69 | S/C | S/C | 72 | ↓ | S/C |
| 49 | ↓ | S/C | 45 | S/C | S/C | 47 | S/C | S/C |
| 42 | ↓ | S/C |  |  |  | 41 | S/C | S/C |
| 37 | ↓ | S/C |  |  |  | 34 | S/C | S/C |
| 31 | ↓ | S/C |  |  |  |  |  |  |
|  |  |  |  | 48 h |  |  |  |  |
| Protein | DENV | LPS | Protein | DENV | LPS | Protein | DENV | LPS |
| 136 | ↓ | S/C | 114 | S/C | ↑ | 250 | ↓ |  |
| 118 | ↓ | S/C | 90 | S/C | ↑ | 123 | S/C | S/C |
| 80 | ↓ | ↓ | 69 | S/C | S/C | 91 | ↓ | S/C |
| 77 | ↓ | ↓ | 64 | ↓ | S/C | 72 | ↓ | S/C |
| 67 | ↑ | ↑ | 52 | ↑ | ↑ | 67 | ↓ | S/C |
| 59 | ↓ | S/C | 45 | ↓ | S/C | 59 | ↓ | S/C |
| 55 | ↓ | S/C | 33 | ↓ | S/C | 55 | ↓ | ↓ |
| 49 | ↓ | S/C |  |  |  | 47 | S/C |  |
| 42 | ↓ | S/C |  |  |  | 42 |  |  |
| 37 |  | S/C |  |  |  | 33 |  |  |
| 31 | ↓ | S/C |  |  |  | 31 |  |  |

### Neuraminidase activity in infected macrophages

Neuraminidase was inhibited using DANA, the transcript was analyzed as described before. Results data were the same proneness to lower residues α-2,3 y α-2,6 (Cross-ref).

Our hypothesis was that neuraminidases are the responsible enzymes to remove sialic acid residues. Data from lectin-blot assays were analyzed by pixel intensity metrics where we do not find significant changes (Cross-ref).

The following procedure was to evaluate the inhibition at the transcriptional level with oseltamivir. The result was a slight decrease of Neu-1 in infected cells at 24 h, while at 48 h there were no significant differences. Infection tends to increase at 24 h regarding infection without oseltamivir, as well as 48 h there are no apparent differences (Cross-ref).

Ultimately, evaluation of enzymatic activity using the fluorophore, which is cleavage by neuraminidase, in the images are observed a higher enzymatic activity at basal levels. When the inhibitor molecule was used, fluorescence fades away and it localizes at the cell surface. Fluorescence was converted to pixel intensity and compared to evaluate the differences between the experimental groups (Cross-ref).

**Figure 1.**
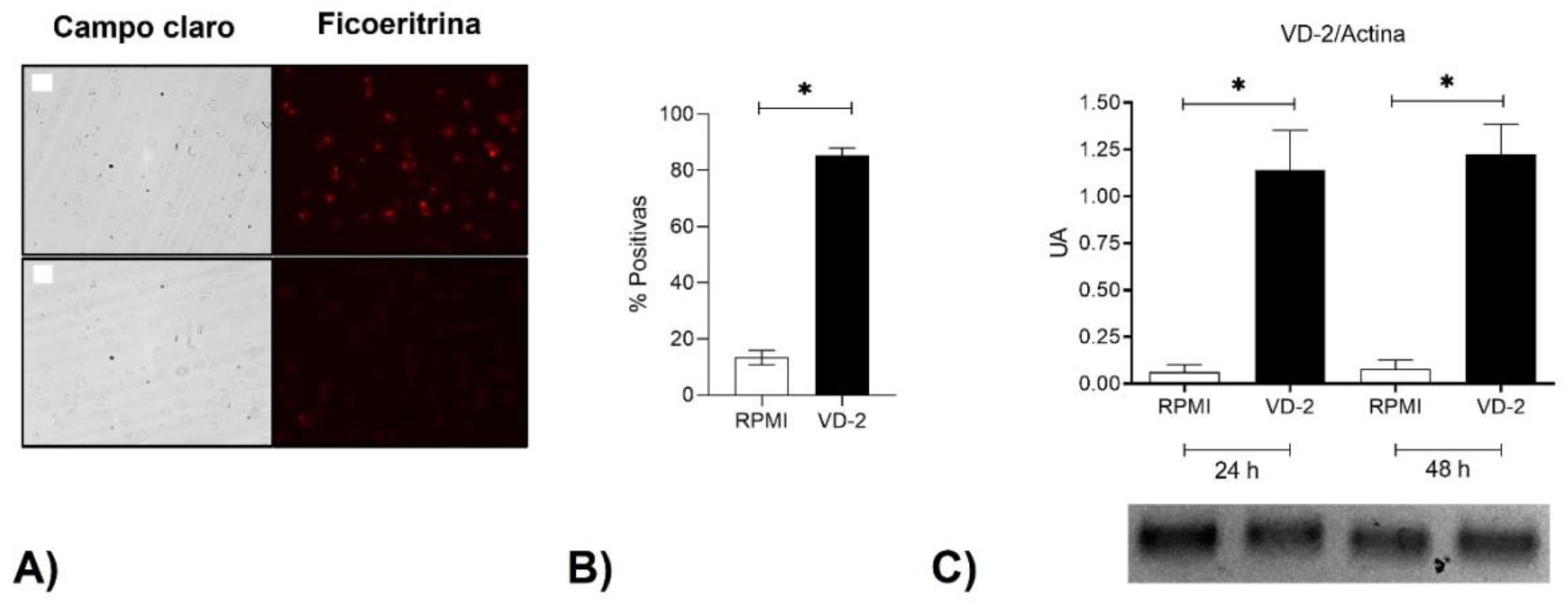
Macrophage’s infection evaluation. A. Top, infected cells. Bottom, non-infected cells. B. Fluorescence positive cells for infection and control treatments at 48 h, at least 200 cells were counted per field. C. Viral genome fragment amplified by RT-PCR in infected and non-infected cells at 24 and 48 h, bands intensity was normalized for pixel intensity according to actin band intensity for each experimental group.

**Figure 2.**
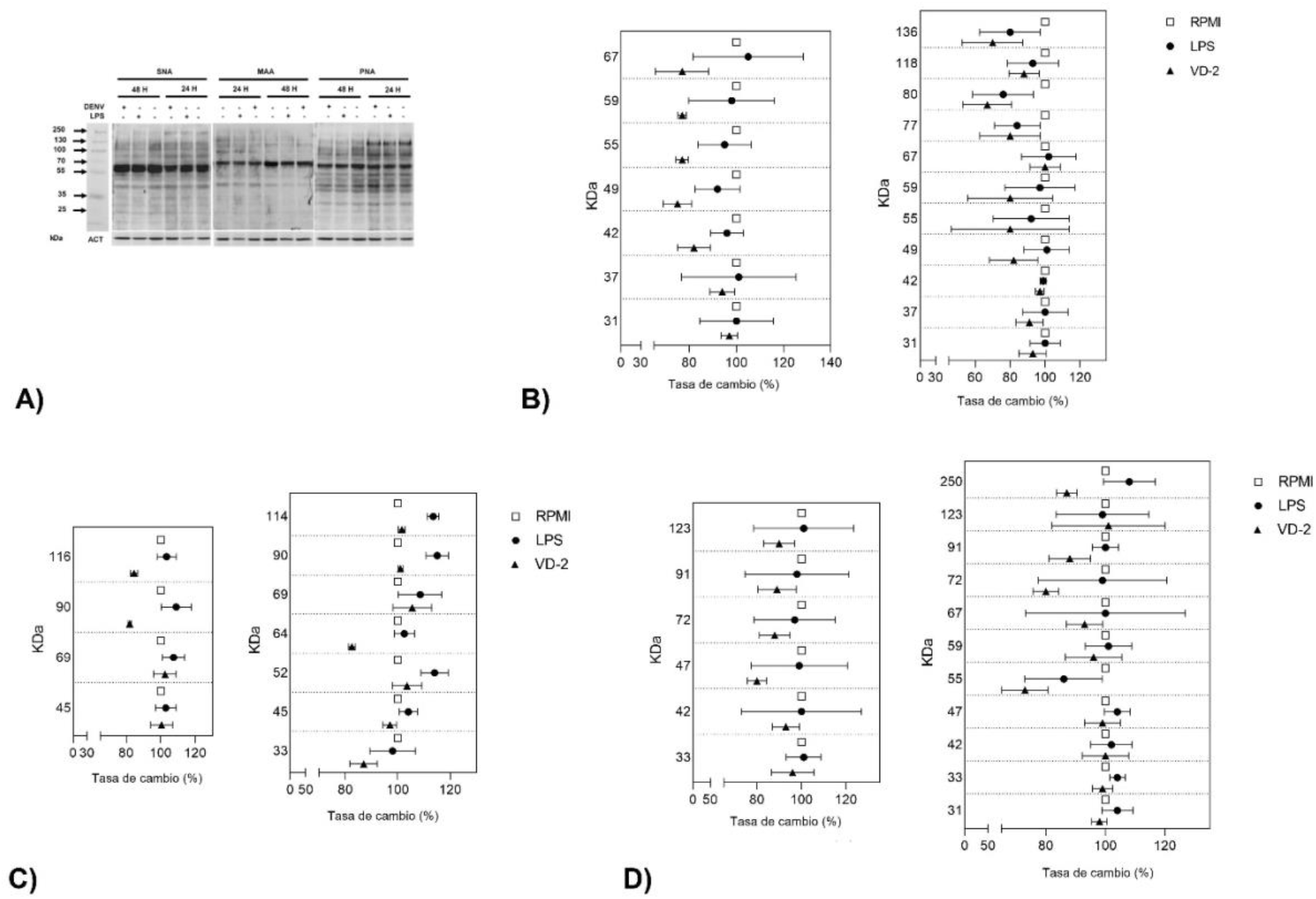
Sialic acid residues content in infected macrophages. A. Total protein reactivity to the three lectins at 24 and 48 h post-treatment. B. Intensity analysis for SNA lectin at 24 and 48 h. C. Intensity analysis for MAA lectin at 24 and 48 h. D. Intensity analysis for PNA lectin at 24 and 48 h.

**Figure 3.**
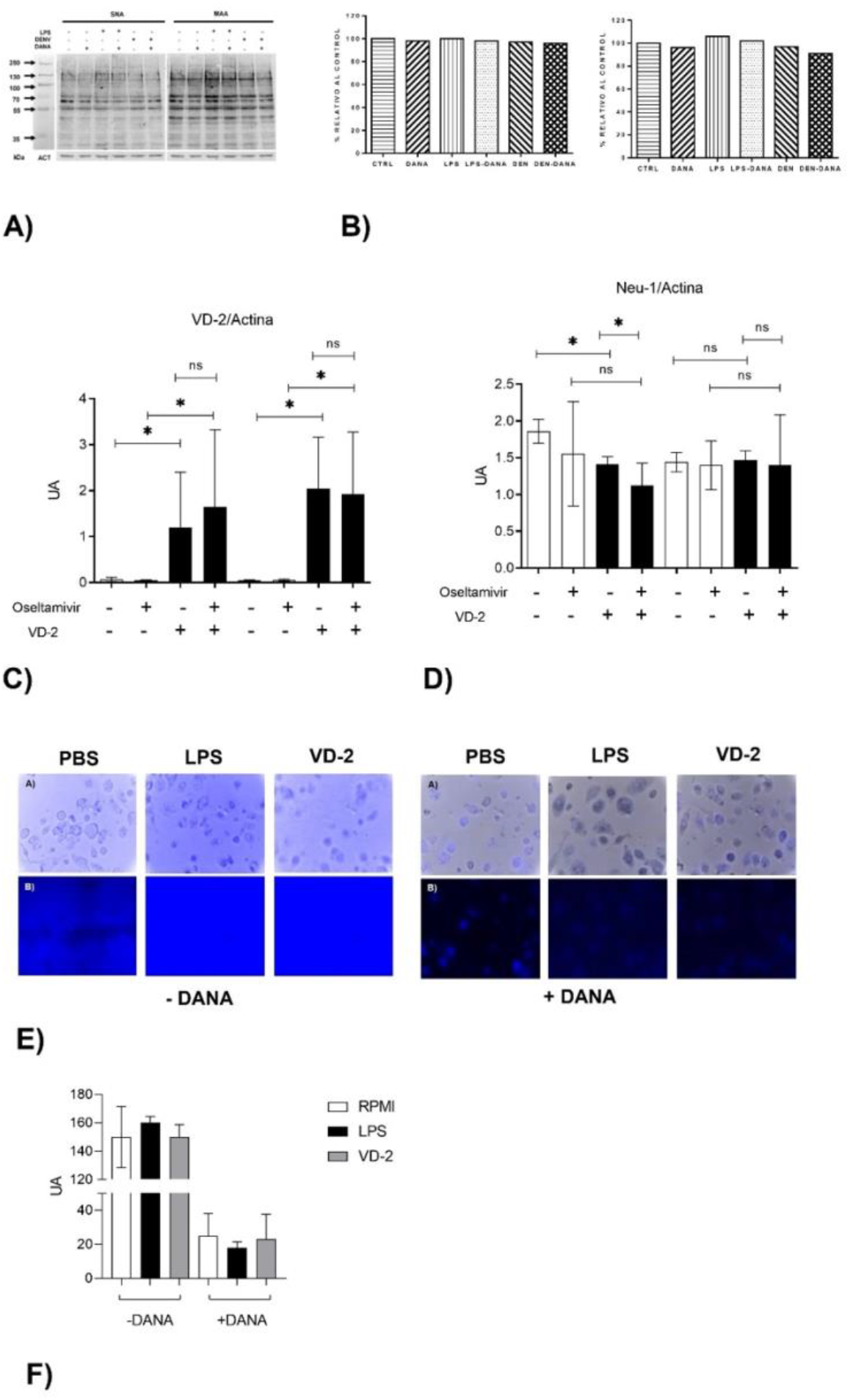
Enzymatic activity for Neu-1 during infection. A. Lectin-blot using previously DANA inhibitor. B. Global changes for SNA (left) and MAA (right) lectins using densitometry analysis. C. Relative infection (RT-PCR) using oseltamivir inhibitor previously. D. Relative Neu-1 expression in macrophages using oseltamivir inhibitor previous viral inoculum. E. Fluorescence assays with DANA and 4-MUNANA substrate. F. Changes analysis in fluorescence assay as intensity pixels.

## Discussion

THP-1 cell line of human monocytes can be artificially differentiated to macrophages using PMA, this process requires the loss of sialic acid residues and a shift in spatial configuration from α-2,3 to α-2,6. This is accompanied by an increase in activity and concentration of neuraminidases in the cell surface, particularly Neu-1 and a lesser proportion of Neu-3 (Stamatos et al. 2005; Liang et al. 2006; Stamatos et al. 2010).

LPS binding to TLR-4 is well detailed in the literature the removal of sialic acid for activation downstream, as today there is no evidence for Dengue infection and sialic acids in macrophages, this is the reason LPS was used as a positive control of proinflammatory profile for monocytes/macrophages (Bonardi et al. 2014; Amith et al. 2010).

In the individual analysis graphics, using SNA at 24 h post-infection, proteins had a decreasing exceptionally to the 31 KDa regarding control, meanwhile for LPS the decreasing tendency is not generalized. At 48 h, there is a pronounced decrease in infected cells with respect to control, in LPS there was a similar fashion. For sialic acid residues α-2,3, only four proteins kept constant in the three treatments, there was a decrease in infection at 24 h, meanwhile in LPS it seems to be an increase for 48 h, while in infection tends to lower, meanwhile to LPS it seems variable.

For β-1,3-galactosamine, at 24 h there is a decrease in all analyzed protein compared to control, regarding LPS there are no notable changes. At 48 h, the sialic acid content is variable, the majority has a decrease and the remaining ones are at an increase.

Some proteins diminish their sialic acid content caused by infection with DENV-2, in SNA lectin, 42, 34, and 31 KDa proteins diminish and also with MAA lectin 33 KDa protein, and their intensity increases with the PNA lectin, which could be a loss of residues of sialic acid which exposes the galactose residues, recognized by β-1,3-gal. Proteins with 67, 59, and 55 KDa besides losing sialic acid, also loses galactoses owing to viral infection, recognition with SNA and PNA decreases, this is due to, loss of in one of them, the other cannot bind to it (Table 1).

Because this decrease was barely visible, we decide to evaluate the transcriptional activity. In the beginning, primers were designed to evaluate Neu-1, −2, −3, and −4 mRNA, but only Neu-1 is present and functional in THP-1 cells differentiated with PMA (Delannoy et al. 2017; Liang et al. 2006).

For determining the neuraminidase enzymatic activities, we used two strategies, the first was to measure the transcript expression using the inhibitor oseltamivir phosphate, insomuch as the molecule inhibits the neuraminidase translocation towards cells surface and further activation, dimerization, and translocation for TLR molecules.

Using the oseltamivir, infection at 24 h showed a slight decrease in Neu-1, this temporary effect could be important but, at 48 h cells, Neu-1 transcription levels were not different to control cells. This is why we suggest the role would be more important in neuraminidase turnover during infection.

The second strategy was to put the cells in contact with the substrate 4-MUNANA with and without the inhibitor DANA, derived from the acetylneuraminic acid, which acts quickly in the sialylation profile over the surface of the cell using non-permeabilized cells.

In the assay with the fluorophore, non-treated cells have a higher enzymatic activity which is confirmed with the DANA inhibitor, on the other hand, infected and LPS-treated cells showed a great neuraminidase activity, fluorescence intensity is saturated, when the inhibitor was used there was no bigger difference regarding control elevated basal activity. The graphic was constructed with pixel intensity per mm^2^, values from infected and stimulated cells were very similar to control, evaluated at 3 min maximum after the substrate was added.

## Conclusions

Macrophages infected with Dengue virus serotype 2 have a globally low-grade decrease in sialic acid content, as individual proteins diminish in a major proportion the sialic acid α-2,6 a las 24 h, meanwhile at 48 h, the more visible changes are in sialic acid α-2,3. Some proteins also lost galactosamine residues, for whatever reason we hypothesize more than one enzyme could be regulating the process. Inhibition at the intracellular level of neuraminidases allows it a slight increase in the infection with Neu-1 transcriptional suppress. Neu-1 basal enzymatic activity in macrophages is high, which to a relative short-times on assays, viral infection or LPS stimulation differences were not measurables with the proposed strategy.

## Author’s contribution

Javier Serrato-Salas: Intellectual authorship. Processing and performance of molecular biology methods. Collection of raw data. Writing and approval of the final version of the manuscript.

Isabel Cruz-Zazueta: Intellectual authorship. Processing and performance of cell culture and SDS-PAGE methods. Collection of raw data. Analysis of results. Writing the initial version of the manuscript.

José Luis Montiel-Hernández: Intellectual authorship. Analysis and interpretation of the results. Critical revision of the manuscript and approval of the final version.

Judith González-Christen: Intellectual authorship and experimental design. Analysis and interpretation of the results. Critical revision of the manuscript and approval of the final version.

